# Mapping the plant extracellular proteome: from fraction-specific secretion patterns to experimental guidelines

**DOI:** 10.64898/2026.11.16.766913

**Authors:** Pengxi Wang, Martin Černý

**Affiliations:** Department of Molecular Biology and Radiobiology, Faculty of AgriSciences, Mendel University in Brno, 613 00 Brno, Czech Republic

**Keywords:** Plant extracellular proteome, sample collection, separation and purification, apoplast, extracellular vesicles, proteomics, method selection

## Abstract

Proteins operating beyond the plasma membrane are central to how plants build and remodel their cell walls, acquire and redistribute nutrients, coordinate developmental processes, and communicate between cells and tissues. Moreover, the reach of this extracellular proteome extends beyond the plant itself, with root and seed exudates shaping the rhizosphere and spermosphere by providing molecular signals and resources that influence microbial communities even before pathogens or symbionts establish direct contact with the plant surface. The extracellular space is therefore a dynamic biological interface connecting processes within the plant to interactions with the soil, microbiome, and broader environment. Despite this central role, the plant extracellular proteome remains fragmented across experimental approaches. Here, we review current techniques for the collection and purification of major plant extracellular fractions and use a uniform prediction pipeline to assess representative publicly available datasets. This analysis reveals consistent fraction-specific patterns in predicted secretion-class composition, with putative unconventional-secretion candidates exceeding canonical candidates in most fractions. We then present a decision tree linking biological questions to appropriate collection and fractionation strategies, together with complementary experimental criteria for assessing extracellular origin that extend beyond sequence-based prediction. Together, these resources provide a framework for interpreting existing extracellular proteomic datasets and for designing future studies that account for the biological distinctness of individual extracellular fractions. They are intended both as an entry point to plant extracellular proteomics and as a reference framework for improving the biological interpretation and comparability of studies across this diverse extracellular landscape.

## Introduction

The release of proteins into the extracellular environment enables organisms to interact with neighboring cells and their surroundings. Animals, plants, fungi and unicellular eukaryotes share an endoplasmic reticulum-Golgi secretory pathway, although their extracellular environments and additional export routes differ; bacteria release proteins through distinct secretion systems and membrane vesicles (Wang et al., 2018; Toyofuku et al., 2023). Plants additionally secrete proteins through unconventional routes that bypass or repurpose the canonical pathway (Ding et al., 2014). The plant cell wall forms a further interface between the plasma membrane and the environment: proteins recovered beyond the membrane may remain soluble in the apoplast, associate with the wall matrix, or occur within extracellular vesicles (EVs) (Rutter and Innes, 2017).

Plant extracellular proteins participate in cell wall remodeling, nutrient transport, intercellular communication, growth and development, and responses to biotic and abiotic stress (Gao et al., 2024; Chen et al., 2025a). Signaling peptides, proteases and cell wall-modifying enzymes further mediate extracellular immunity and defense against pathogens (Jian et al., 2024). For example, leaf apoplast proteomics has revealed changes in extracellular protein composition during cold acclimation in jojoba (Zheng et al., 2026) and *Ammopiptanthus mongolicus* (Zheng et al., 2023), and pattern-triggered immunity alters the abundance of defense-related proteins in the *Arabidopsis* leaf apoplast (Chen et al., 2025).

Proteins can reach these extracellular samples through several routes: canonical secretion, unconventional secretion, developmentally programmed cell death, and leakage from damaged cells. The protein composition of sequentially collected xylem sap fractions may vary following stem cutting (Kopecká and Černý, 2024). Developmentally regulated cell death in the Arabidopsis root cap may contribute intracellular proteins to extracellular samples (Wang et al., 2024). Such observations show that the sample collected and the fraction ultimately analyzed are not necessarily equivalent: apoplastic washing fluid, wall-bound extracts, EV preparations and exudates each recover different protein populations (Fig. 1). We therefore use extracellular proteome for the proteins recovered from a given sample, and secretome for proteins actively released from cells through secretion pathways. These terms are not necessarily interchangeable because extracellular samples may also contain proteins released through cellular damage or leakage.

**Fig. 1.**
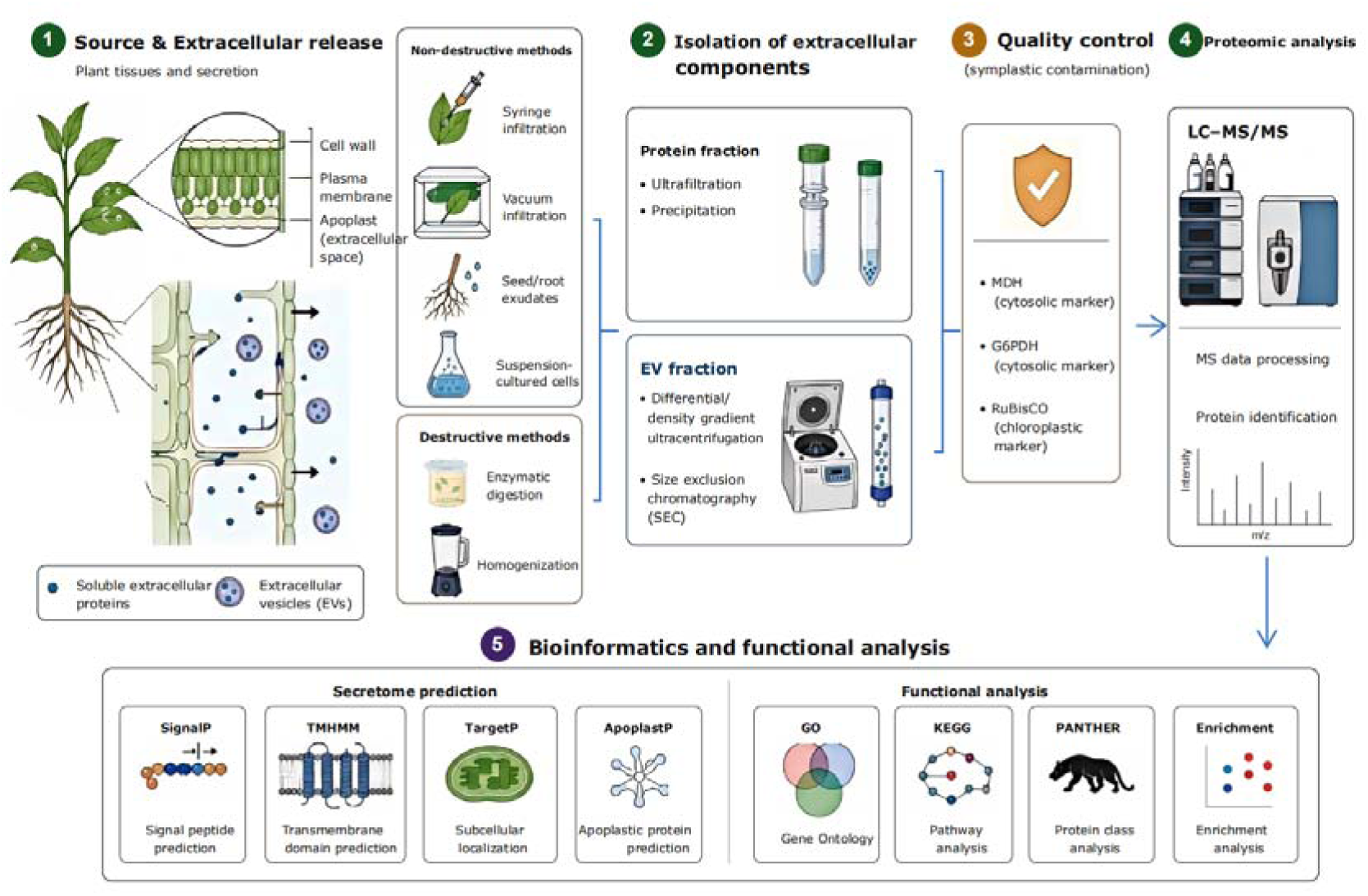
Workflow for extracellular proteomic profiling in plants.

Recent advances in plant extracellular vesicle markers, isolation methods, and characterization practices, together with the increasing availability of public proteomics datasets, have created opportunities for comparative analysis (Perez-Riverol et al., 2022; Thieron et al., 2024; Rodríguez de Lope et al., 2025). Nevertheless, the interpretation of extracellular proteomic data depends on both the starting sample and the fraction ultimately analyzed. Unfractionated leaf apoplastic washing fluid, its soluble fraction, and an EV-enriched fraction cannot be treated as equivalent samples. Moreover, most proteins identified in an Arabidopsis leaf EV preparation lacked a predicted classical signal peptide, illustrating why signal-peptide prediction alone cannot establish how proteins reached the extracellular space (Rutter and Innes, 2017). Differences in sample sources, collection and isolation procedures, and protein secretion and localization prediction tools complicate comparisons across studies. Whether common patterns in predicted protein classes emerge across datasets obtained from different plant materials and isolation procedures therefore remains an important question.

This review compares methods for collecting plant extracellular samples and separating their protein-containing fractions. The numbers and relative abundances of predicted protein classes are then examined across nine proteomic datasets. Based on these findings and published methodological evidence, a decision tree is developed to guide method selection. We then discuss proteomics-based strategies for evaluating extracellular association, quality control, and the limits of inferring protein release mechanisms.

## 1. Biological origins of proteins recovered from plant extracellular samples

### Secretory routes across kingdoms and in plants

Protein secretion occurs across all kingdoms of life through both conserved and lineage-specific mechanisms. Animals, plants, fungi, and other unicellular eukaryotes share the conventional endoplasmic reticulum (ER)-Golgi secretory pathway but also employ unconventional protein secretion (UPS) routes and release extracellular vesicles, although their extracellular environments and additional export routes differ (De Vallée et al., 2023; Pope and Prade, 2026). Plant and fungal cell walls also affect protein recovery. Bacteria instead use Sec/Tat translocation pathways, specialized secretion systems, and membrane vesicles to export proteins (Toyofuku et al., 2023).

In plants, the canonical secretory pathway typically begins with an N-terminal signal peptide directing a protein into the ER, followed by trafficking through the Golgi apparatus and trans-Golgi network toward the cell surface. UPS encompasses routes that deviate from this pathway. Four UPS routes have been proposed: Golgi bypass (type I), vacuole-mediated release (type II), multivesicular body (MVB)-mediated release (type III), and exocyst-positive organelle (EXPO)-mediated release (type IV) (Ding et al., 2014). Importantly, these routes describe potential transport mechanisms rather than protein classes that can be assigned from sequence alone. The absence of a predicted signal peptide therefore does not establish that a protein was released through UPS (Robinson et al., 2016).

MVBs and EXPOs illustrate the diversity of vesicle-associated protein export in plants. MVBs contain intraluminal vesicles that can be released into the apoplast following fusion with the plasma membrane. In *Arabidopsis*, EVs have been implicated in the transfer of small RNAs to the fungal pathogen *Botrytis cinerea* (Cai et al., 2018). More recent work showed that TET8- and PEN1-marked EVs form largely distinct populations whose release responds to biotic stress (Koch et al., 2025). The functional importance of EV protein cargo was further demonstrated in rice, where EV-associated defense proteins were transferred into *Rhizoctonia solani* and reduced disease susceptibility (Huang et al., 2025). EXPO, first characterized as a double-membrane organelle involved in cytosol-to-cell-wall export, represents another vesicle-associated route (Wang et al., 2010). Recent structural and functional analyses identified Exo70E2-positive EVs derived from EXPO and linked this pathway to plant defense (Gao et al., 2026). Together, these studies demonstrate the diversity of plant EV populations and their possible biological origins. However, identifying a protein in an EV-enriched fraction does not establish its individual export route, particularly because EV preparations may also contain co-isolated non-vesicular material.

### Other origins of proteins in collected samples

In addition to canonical and unconventional secretion, proteins may enter extracellular samples through natural cell detachment, developmentally programmed cell death, or cellular damage during sample collection and processing (Charura et al., 2024). Root border cells naturally detach from the root cap and contribute intracellular and extracellular material to the surrounding environment (Ropitaux et al., 2020; Kranawetter and Sumner, 2025). Distinct from cell detachment, programmed cell death can also release cellular contents. Studies in *Arabidopsis* have shown that root cap cell death and the subsequent clearance of cellular remains are regulated developmental processes associated with microbial colonization (Charura et al., 2024; Wang et al., 2024). Developmental timing is also relevant to seed samples. In barley, aleurone cells secrete hydrolytic enzymes during germination and subsequently undergo hormonally regulated programmed cell death (Domínguez and Cejudo, 2014). These processes may contribute intracellular proteins to extracellular samples, although their contribution to individual proteomic datasets requires independent evidence.

Sample collection and processing may introduce intracellular proteins through cellular damage. Tissue homogenization, stem cutting for xylem sap collection, leaf infiltration, and handling of germinating seeds may disrupt cells and release their contents. Kopecká and Černý (2024) examined sequentially collected xylem sap fractions, highlighting the importance of collection procedures when interpreting recovered proteins. However, proteins predicted to have intracellular localization cannot automatically be attributed to sampling damage, as they may also arise from regulated release, developmental processes, or incompletely characterized export pathways. Protein origin must therefore be interpreted in the context of sample collection, tissue integrity, and independent localization or functional evidence.

## 2. Collection of plant samples for extracellular proteomics

The choice of sampling method determines which extracellular proteins can be collected before any protein extraction or purification begins. Four targets guide sampling: soluble apoplastic proteins, cell wall-bound proteins, EV-associated proteins, and proteins released into the environment surrounding the plant (Rutter and Innes, 2017). These targets are not separate anatomical spaces. For example, apoplastic washing fluid (AWF) contains soluble proteins and may also contain cell wall proteins and EVs, whereas a root exudate can contain free proteins, vesicles, and material released with root-cap cells (Kranawetter and Sumner, 2025).

### Soluble apoplastic proteins

Infiltration followed by low-speed centrifugation is the principal approach for collecting leaf AWF. Water or a defined buffer is introduced into the intercellular space by vacuum or syringe infiltration; fluid is then recovered from the tissue by centrifugation. Recent Arabidopsis studies have used this approach to profile the leaf apoplast during immune responses and to examine the effects of collection conditions on protein recovery and cytosolic leakage (Chen et al., 2025a; Chen, Karinshak and Vierstra, 2025b). Syringe infiltration is useful for individual leaves or small experiments, while vacuum infiltration can process more tissue in parallel. Gravity-based collection and water displacement have also been applied to leaves or stems, but are less established across species (Pechanova et al., 2010; Wu et al., 2014).

Methods should be compared in terms of fluid recovery per unit tissue mass and the composition of the recovered material, rather than assuming that any method yields an inherently pure extracellular sample. Infiltration dilutes the native apoplastic fluid, while excessive infiltration pressure or centrifugal force may damage cells and introduce intracellular material into the recovered fluid. Tissue age, buffer composition, infiltration time, and centrifugal force should therefore be reported to facilitate comparisons across studies. AWF is a starting sample rather than an isolated soluble-protein fraction, because soluble proteins and EVs can be separated during subsequent processing (Rutter and Innes, 2017; Eldahshoury et al., 2024). For example, soluble apoplastic proteins were recovered as the supernatant after ultracentrifugation of the total apoplastic extract at 200,000 × g for 20 min at 4 °C (Lee et al., 2025).

### Cell wall bound proteins

Cell wall bound proteins require a different sampling decision because many remain associated with the wall after soluble AWF has been removed. One route begins with intact leaves, stems, or roots and exposes accessible wall proteins to an infiltration solution. The other begins with harvested tissue that will undergo disruption and preparation of a wall-enriched fraction. A comparison of sugarcane stem methods showed that non-destructive and destructive approaches recovered different sets of cell wall proteins (Calderan-Rodrigues et al., 2016). Recent studies have also sampled roots and leaves for cell wall proteomics under stress conditions (Chen et al., 2022).

For cell wall-associated proteins, sampling records should specify the organ, developmental stage, fresh mass, and tissue integrity before collection. Unlike AWF collection, preparation of a wall-enriched fraction typically involves tissue disruption followed by separation of cell wall material. Salt extraction may then be applied either to intact tissue or to a prepared wall fraction, depending on the experimental workflow. Distinguishing these approaches is important because proteins extracted from accessible wall surfaces and those recovered from wall-enriched preparations may represent different protein populations (Albenne et al., 2013).

### EV associated proteins

Plant EV studies first collect a fluid in which vesicles may be present; the vesicles are then isolated from that fluid. Leaf AWF is the best developed starting material and has been used for EV proteomics in Arabidopsis (Rodríguez de Lope et al., 2025). Other reported sources include root exudates, conditioned culture medium, and phloem sap. EVs have been recovered from tomato root exudates, Arabidopsis cell cultures, and melon phloem sap collected after stem incision (De Palma et al., 2020; Yugay et al., 2023; Sánchez-López et al., 2024).

The source fluid and its collection procedure should be specified before describing the EV isolation method. AWF, root exudates, culture medium, and vascular sap represent different extracellular environments and may contain distinct non-vesicular components. Tissue homogenates and expressed fruit juices require particular caution because mechanical disruption can release cellular debris and intracellular vesicles, as well as generate vesicle-like particles during processing (Huang et al., 2025; Yang et al., 2025). Consequently, the origin of vesicles recovered from disrupted tissues cannot be determined by size alone. The choice of starting material and collection procedure therefore influences the composition and biological interpretation of the isolated vesicle population.

### Proteins released outside the plant and other accessible fluids

Root and seed exudates are collected from the surrounding medium without extracting an internal tissue fraction. Root exudates can be obtained from plants grown in hydroponic systems or transferred to a defined collection solution. This approach permits timed sampling, but plant transfer, collection-medium composition, sampling duration, and microbial activity can influence the recovered material (McLaughlin et al., 2025). In Arabidopsis, activation of stress-related mitogen-activated protein kinases altered the proteins detected in root exudates (Strehmel et al., 2017). Root-tip mucilage can also be collected separately from bulk root exudates; a wheat study analyzed mucilage and soluble root-secreted proteins as distinct preparations (Staudinger et al., 2022). Seed exudates are collected from the surrounding medium during seed imbibition. Their composition may vary with seed integrity, imbibition duration, germination stage, and collection volume, making these parameters important to report (Tiddia et al., 2023).

Xylem and phloem sap also yield proteins, although they are internal transport fluids before collection (Carella et al., 2016). Xylem sap can be collected from a cut stem under root pressure or with an applied-pressure method; recent proteomic work has used xylem sap to examine root–shoot responses to flg22 (Kopecká and Černý, 2024). Phloem sampling includes EDTA-facilitated exudation from excised leaves, incision-derived sap in suitable species, and aphid stylectomy (Broussard et al., 2023). An Arabidopsis proteomic study used EDTA-facilitated collection, while a melon study obtained phloem sap by stem incision (Sánchez-López et al., 2024; Moreno-Sanguino et al., 2026). Sap emerging from a wound is thus a valid sampling route, but it may contain material released from damaged cells and adjacent tissues (Broussard et al., 2023). Wound exudate should specify the tissue cut, the fluid collected, and the collection interval rather than imply a single biological source.

Conditioned medium from suspension-cell cultures provides another accessible source. Cells can be removed from the medium after a defined culture or treatment period, as demonstrated in secretome studies of sorghum cells under heat stress (Ngcala et al., 2020) and Pinus nigra embryogenic cultures (Pernis et al., 2026). Culture conditions and cell viability should be documented to assess potential intracellular contamination. However, culture-medium proteomes may not fully represent the extracellular environment of intact plant organs.

Specialized plant surfaces offer further sampling routes. Exudate from a wet stigma can be collected as visible droplets or by gentle surface sampling; proteomic work on lily and olive used these approaches to study the stigma secretion itself, rather than a whole stigma extract (Rejón et al., 2013). Floral and extrafloral nectar can be collected from nectaries with a capillary or syringe, and both have supported protein identification (Silva et al., 2020; Nascimento et al., 2018). Guttation droplets are accessible at leaf margins, but current sampling studies mainly address their metabolites and physiology; their value for broad extracellular protein profiling remains less established (Routaboul et al., 2024). These surface fluids should be sampled at a defined developmental stage and protected from pollen, microbes, and environmental deposits when the aim is to study plant proteins.

## 3. Separation and purification of plant extracellular proteins

The samples described in Section 2 are starting materials for protein preparation. Their proteins may be dissolved in a collected fluid, associated with cell-wall material, or carried by EVs. Each form requires different processing. In particular, concentrating an entire AWF sample does not separate its soluble proteins from vesicle-associated proteins. The intended fraction must therefore be defined before a purification method is selected.

### Proteins in extracellular fluids and exudates

Extracellular fluids and exudates can be processed to recover proteins from the collected sample or to isolate a soluble-protein fraction after particle removal. AWF, root and seed exudates, xylem sap, and cell-culture medium often require concentration before proteomic analysis. Centrifugal ultrafiltration concentrates material above a selected molecular-weight cutoff and permits buffer exchange. For example, *Arabidopsis* AWF was concentrated using a 3-kDa filter before protein quantification and peptide preparation (Chen et al., 2025a). In cotton fibers, the total apoplastic fraction was separated by ultracentrifugation at 200,000 × g for 20 min into a soluble supernatant and a particle-enriched pellet (Lee et al., 2025). When soluble proteins are the target, particles should be removed before concentration, because ultrafiltration alone does not separate soluble proteins from particles (Eldahshoury et al., 2024).

Once the target fluid fraction has been selected, lyophilization and protein precipitation provide alternative methods for recovering proteins from the whole collected fluid or its soluble supernatant. Arabidopsis root-exudate medium was filtered and lyophilized before protein digestion (Strehmel et al., 2017). Proteins from cell-free sorghum suspension-culture medium were recovered by acetone precipitation, followed by pellet solubilization (Ngcala et al., 2020). Xylem sap was lyophilized, washed with organic solvents, and processed for protein digestion and peptide desalting (Kopecká and Černý, 2024).

These preparation methods can influence the proteins ultimately detected. During ultrafiltration, small proteins and peptides may pass into the filtrate, depending on the molecular weight cutoff. For example, a 10-kDa filter was used to obtain separate phloem-exudate fractions enriched in larger proteins and smaller proteins or peptides, and both fractions were analyzed (Moreno-Sanguino et al., 2026). Precipitation and solvent washing may also result in protein loss. Therefore, studies should report both the preparation method and the fraction used for proteomic analysis, because different procedures may recover different proteins.

### Cell wall associated proteins

Recovering proteins retained in the cell wall matrix generally requires extraction beyond fluid concentration. A common approach first enriches a cell wall fraction from disrupted tissue and washes it to reduce soluble material. Proteins are then released from the isolated wall fraction with salt solutions. CaCl and LiCl were used to extract proteins from isolated cell wall fractions of Brachypodium distachyon (Douché et al., 2021). Sequential CaCl and LiCl extraction was used to recover proteins held by weak or ionic interactions in Marchantia polymorpha (Kolkas et al., 2022). The resulting extract is thus a defined salt extractable wall protein fraction, not the complete wall proteome. Proteins that remain tightly attached may need a different approach, such as digestion of the isolated wall fraction (Kolkas et al., 2022). Because cell wall preparation involves tissue disruption, intracellular carryover remains an interpretation issue even after extensive washing. Cell wall proteomics reviews likewise emphasize how extraction and wall preparation affect the recovered proteins (Calderan-Rodrigues et al., 2019).

### EV associated proteins

For studies of EVs present outside intact plant cells, the source of the vesicles matters as much as the separation method. AWF or conditioned medium can contain vesicles released into an extracellular fluid (Rutter and Innes, 2017; Yugay et al., 2023). Tissue homogenates can also yield membrane particles, but their isolation does not establish that those particles existed extracellularly before disruption. These preparations should not be treated as equivalent when interpreting an EV proteome (Thieron et al., 2024; Rodríguez de Lope et al., 2025).

Plant AWF studies commonly begin by removing cells and debris, followed by differential centrifugation to obtain EV-enriched fractions (Rutter and Innes, 2017; Gao et al., 2024). Successive centrifugation steps remove different particle populations, while the final high-speed pellet may contain EVs together with other sedimentable material (Huang et al., 2021; Welsh et al., 2024). Successive centrifugation at 2,000, 10,000, and 25,000 × g was used before small EVs were enriched from *Nicotiana benthamiana* AWF at 100,000 × g (Eldahshoury et al., 2024). EV fractions obtained from *Arabidopsis* AWF at different centrifugal forces were also compared and further separated using density gradients (Huang et al., 2021). The resulting pellet should therefore be described as an EV-enriched fraction rather than a pure EV preparation (Welsh et al., 2024).

Further purification can reduce overlap between EVs and other components of the starting fluid (Thieron et al., 2024). Density-gradient centrifugation separates particles by buoyant density, whereas size-exclusion chromatography (SEC) separates larger particles from smaller soluble components as they pass through a porous column (Liangsupree et al., 2021). Ultrafiltration can concentrate a particle-containing fluid before further separation, but does not alone establish that the retained particles are EVs. Other particles and macromolecules may also be retained, and membrane clogging can affect recovery (Liangsupree et al., 2021). Ultrafiltration followed by SEC was used to prepare EV fractions from *Arabidopsis* and *Brassica oleracea* AWF for proteomic marker analysis (Rodríguez de Lope et al., 2025). These methods therefore serve different functions within a separation workflow, and their performance should be considered in relation to the starting material and the fraction selected for analysis (Thieron et al., 2024; Welsh et al., 2024).

Other approaches, including polymer precipitation, aqueous two-phase separation, immunoaffinity capture, and microfluidic isolation, may also be considered for EV recovery (Weng et al., 2016; Kırbaş et al., 2019; Jia et al., 2022). Polymer precipitation may co-recover non-vesicular macromolecules, whereas immunoaffinity capture selects vesicles bearing the targeted surface marker (Liangsupree et al., 2021; Jia et al., 2022). Their applicability to plant extracellular fluids requires evidence from comparable samples (Thieron et al., 2024). Results obtained from animal biofluids or disrupted plant tissues cannot, by themselves, establish their performance for EVs collected from plant AWF or conditioned medium.

After an EV-enriched fraction has been obtained, its proteins can be extracted for proteomic analysis (Gao et al., 2024). Detection in that fraction establishes association with the preparation, but does not distinguish proteins enclosed within vesicles from those attached to vesicle surfaces or co-isolated during separation (Welsh et al., 2024). Particle measurements, microscopy, protein markers, and cargo-protection assays provide complementary evidence for characterizing the preparation and assessing protein localization. Plant EV marker analysis provides a basis for evaluating EV-enriched proteomes (Rodríguez de Lope et al., 2025), while reporting guidance emphasizes distinguishing what each separation and characterization step demonstrates from what remains uncertain (Welsh et al., 2024).

## 4. Comparative analysis of plant extracellular protein fractions

We reanalyzed nine plant extracellular proteomic datasets using six prediction tools: SignalP 6.0, DeepTMHMM 1.0, LSPpred, OutCyte, DeepLoc 2.1 and ApoplastP (Table 1). The datasets included suspension-cell culture medium, seed and root exudates, leaf apoplastic washing fluid (AWF), xylem and phloem sap, an AWF-derived EV fraction, a soluble cotton fibre apoplastic fraction, and a cell-wall protein preparation. Proteins were assigned to four prediction categories using the same criteria in all nine datasets: canonical secretion candidate (C), unconventional secretion candidate (U), intracellular-like (I), and other/unresolved (O). Canonical candidates required a predicted signal peptide without an additional transmembrane helix. Among proteins with neither feature, unconventional candidates required a high-confidence LSPpred “LSP” prediction; intracellular-like candidates required LSPpred “Non-LSP”, OutCyte “Intracellular”, a DeepLoc prediction other than “Extracellular”, and ApoplastP “Non-apoplastic”. Proteins meeting none of these criteria were assigned to the other/unresolved class (Table 2; Fig. 2). Because the original studies used different collection and fractionation procedures, we compared the protein compositions of the fractions analyzed, rather than treating all nine datasets as equivalent samples of the extracellular proteome.

**Fig. 2.**
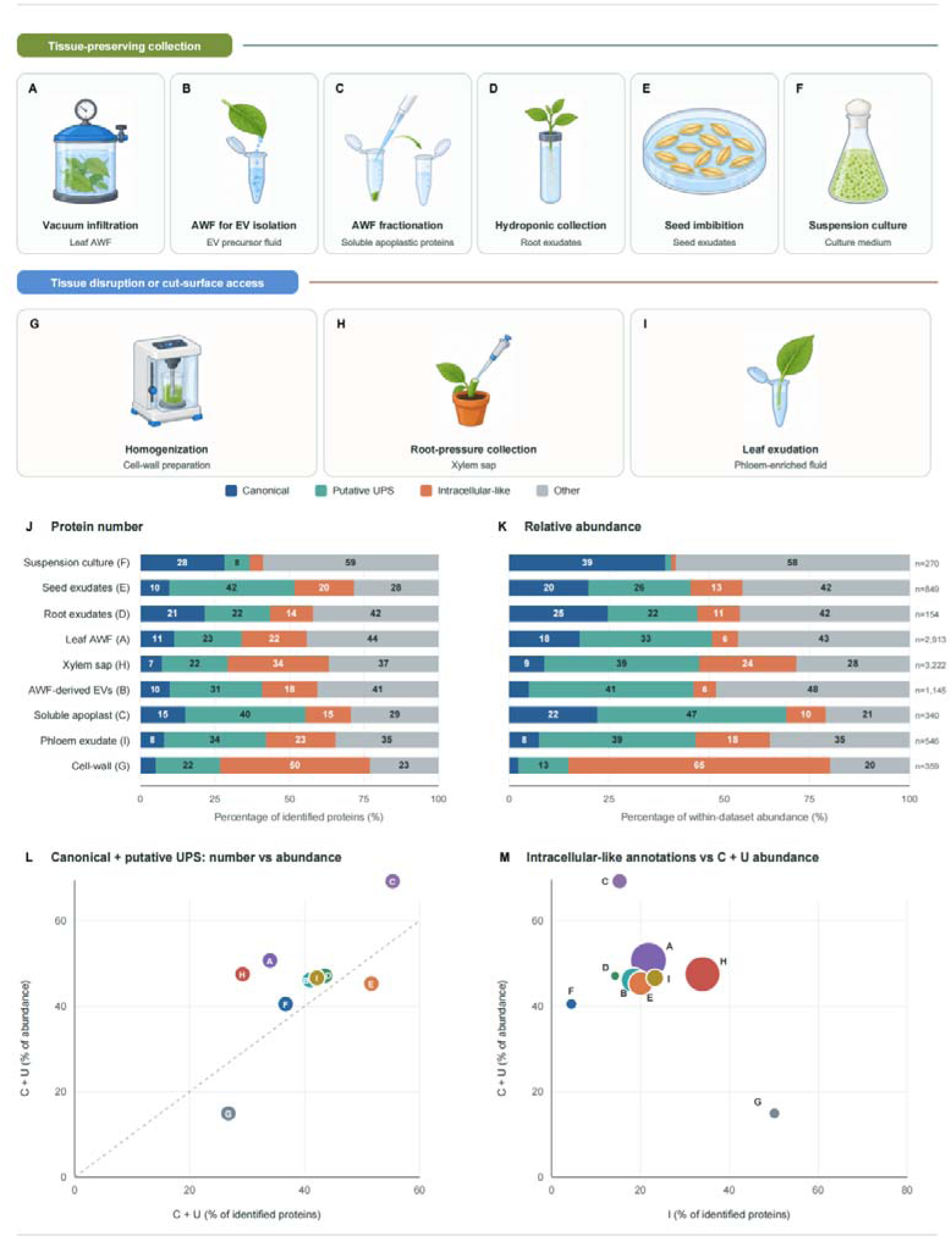
Plant extracellular protein sampling and predicted protein-class composition across nine datasets. (A– I) Collection routes and analyzed samples or fractions: leaf apoplastic washing fluid (AWF; A), AWF used for extracellular vesicle (EV) isolation (B), soluble apoplastic fraction obtained from AWF (C), root exudates (D), seed exudates (E), suspension-cell culture medium (F), cell-wall protein preparation (G), xylem sap (H), and phloem-enriched fluid (I). (J, K) The percentage of identified proteins (J) and the percentage of total protein abundance (K) assigned to canonical secretion candidates (C), unconventional secretion candidates (U), intracellular-like predictions (I), or unresolved proteins (O) within each dataset. Values within bars are rounded; n is the number of identified proteins. (L) The percentage of identified proteins classified as C + U plotted against their percentage of total protein abundance; the dashed line indicates equal percentages. (M) The percentage of identified proteins classified as I plotted against the percentage of total protein abundance classified as C + U; bubble area is proportional to the number of C + U proteins. Letters in L and M refer to A– I. The classes are predictions: the percentage classified as I is not a measured contamination rate. Percentages describe each analyzed dataset, not protein yield across methods. Exact values and dataset sources are given in Table 2.

**Table 1.** Bioinformatic tools and resources relevant to plant extracellular protein analysis.

| Tool /resource | Function | Website | Status | Reference |
|---|---|---|---|---|
| Phobius | Signal peptides and transmembrane topology | https://phobius.sbc.su.se/ | Active | Kall et al.,2004 |
| iPSORT | N-terminal protein sorting signals | https://ipsort.hgc.jp/ | Available | Bannai et al.,2002 |
| Plant-mPLoc | Plant protein subcellular localization | https:/www.csbio.sjtu.edu.cn/bioinf/plant-muli/ | Active | Chou and Shen,2010 |
| Predotar | N-terminal organelle targeting sequences | https://urgi.versailles.inra.fr/Tools/Predotar | Available | Smallet al.,2004 |
| PTS1 predictor | Peroxisomal targeting signal 1 | https://mendel.imp.ac.at/pts1/ | Active | Neuberger et al.,2003 |
| TargetP2.0 | N-terminal secretory and organelle targeting signals | https:/services.healthtech.dtu.dk/services/TargetP-2.0/ | Active | Almagro Armenteros et al.,2019 |
| WoLF PSORT | Eukaryotic protein subcellular localization | https://wolfpsort.hgc.jp/ | Available | Horton et al.,2007 |
| Plant Cell Marker | Plant cell marker database | https://www.tobaccodb.org/pcmdb/ | Active | Jin et al.,2022 |
| SUBA5 | Arabidopsis subcellular localization database | https://suba.live/ | Active | Hooper et al.,2022 |
| cropPAL2020 | Crop protein subcellular localization resource | https://croppal.org/ | Active | Hooper et al.,2020 |
| LSPpred | Plant leaderless secretory protein prediction | http:/1sppred.Ispdb.org/ | Available | Lonsdale et al.,2023 |
| Cell wall genomics | Plant cellwall resource links | https://cellwall.genomics.purdue.edu/inkslindex.html | Available | Website only |
| CAZy | Carbohydrate-active enzyme families and functions | https://ww.cazy.org/ | Active | Drula et al.,2022 |
| DeepTMHMM 1.0 | Transmembrane protein topology | https://services.healthtech.dtu.dk/services/DeepTMHMM-1.0/ | Active | Hallgren et al.,2022 |
| ApoplastP | Apoplastic localization of plant proteins | https://apoplastp.csiro.au/ | Available | Sperschneider et al.,2018 |
| WallProtDB-2 | Plant cell wall proteome database | https://www.polebio.Irsv.ups-tlse.frallProtDB/ | Available | San Clemente et al.,2022 |
| SignalP 6.0 | Signal peptides and cleavage stes | https://services.healthtech.dtu.dk/services/SignalP-6.0/ | Active | Teufel et al.,2022 |
| DeepLoc 2.1 | Subcellular localization and membrane association | https://services.healthtech.dtu.dk/services/DeepLoc-2.1/ | Active | Odum et al.,2024 |
| OutCyte | Putative unconventional protein secretion | https://www.outcyte.com/ | Active | Zhao et al.,2019 |
| BUSCA | Integrated subcellular localization prediction | http://busca.biocomp.unibo.it | Active | Savojardo et al.,2018 |
| PROSITE | Protein families,domains and functional-site motifs | https://prosite.expasy.org/ | Active | Sigrist et al.,2026 |

**Table 2.** Plant extracellular protein sampling and class composition.

| **Sample/dataset** | **Sample collection**<br>**(Section 2)** | **Separation /purification**<br>**(Section 3);analyzed fraction** | **N** | **Canonical**<br>**(C)** | **Putative UPS**<br>**(U)** | **Intracellular-like**<br>**(1)** | **Other**<br>**(0)** |
|---|---|---|---|---|---|---|---|
| **Suspension-cell culture medium**<br>**PXD021536** | Sorghum suspension-culture medium | Fiter out cells;acetone-precipitate medium proteins.<br>**Analyzed:**culture-medium extract. | 270 | 76<br>28.1% \| 39.0% | 23<br>8.5% \| 1.5% | 12<br>4.4% \| 1.1% | 159<br>58.9% \| 58.4% |
| **Seed exudates**<br>**PXD081462** | 10 barley seeds in 1mL water; collect liquid after 1-4h imbibition | Lyophilize and process using the authors' methanol-based procedure.<br>**Analyzed:**seed-exudate proteins. | 849 | 83<br>9.8% \| 19.8% | 355<br>41.8% \| 25.5% | 170<br>20.0% \| 13.0% | 241<br>28.4% \| 41.7% |
| **Root exudates**<br>**PXD006328** | Liquid collected around hydroponic Arabidopsis roots | Filter (0.45µm),lyophilize,and prepare proteins.<br>**Analyzed:**root-exudate proteins. | 154 | 33<br>21.4% \| 24.7% | 34<br>22.1% \| 22.4% | 22<br>14.3% \| 10.6% | 65<br>42.2% \| 42.4% |
| **Leaf AWF**<br>**PXD060654** | Arabidopsis leaf AWF;vacuum infiltration and low-speed centrifugation | Concentrate with a 3-kDa filter.<br>**Analyzed:**AWF proteins,without EV/soluble separation. | 2,913 | 332<br>11.4%117.6% | 657<br>22.6% \| 33.1% | 635<br>21.8% \| 6.5% | 1,289<br>44.2%142.8% |
| **Xylem sap**<br>**PXD053406** | Sap collected in successive fractions after stem cutting | Flash-freeze,lyophilize,solvent-wash,and solubilize.<br>**Analyzed:**xylem-sap proteins. | 3,222 | 233<br>7.2% \| 8.8% | 708<br>22.0% \| 38.7% | 1,094<br>34.0% \| 24.2% | 1,187<br>36.8% \| 28.3% |
| **AWF-derived EVs**<br>**PXD058098** | Arabidopsis leaf AWF | Clarify,ultrafilter,separate by SEC,and extract EV proteins.<br>**Analyzed:**EV-enriched fraction. | 1,145 | 112<br>9.8% \| 4.9% | 356<br>31.1% \| 41.2% | 211<br>18.4% \| 5.6% | 466<br>40.7%148.3% |
| **Solubleapoplastic fraction**<br>**PXD051721** | 9-DPA Gossypium hirsutum fibers;total apoplastic wash fluid | Centrifuge at 200,000×g for 20 min;collect the supernatant and acetone-precipitate proteins<br>**Analyzed:**APO_s200 soluble fraction. | 340 | 51<br>15.0% \| 22.0% | 137<br>40.3% \| 47.2% | 52<br>15.3% \| 9.8% | 100<br>29.4% \| 21.0% |
| **Phloem exudate**<br>**PXD065325** | Exudate collected from cut Arabidopsis leaves | Separate with a 10-kDa fiter.<br>**Analyzed:**combined lists from retained fraction A and fitrate C. | 546 | 43<br>7.9% \| 7.5% | 187<br>34.2% \| 39.1% | 127<br>23.3% \| 18.5% | 189<br>34.6% \| 34.9% |
| Cell-wall protein<br>PXD012761 | Sugarcane leaf or internode tissue | Extract cellwall roteins.<br>**Analyzed:**wall-enriched protein preparation | 359 | 18<br>5.0% \| 2.3% | 78<br>21.7% \| 12.6% | 180<br>50.1% \| 65.2% | 83<br>23.1% \| 19.9% |

Clasication:C=SignalP -posive,no adifional TM;U=no signal peptide or TM and high-cnfience LSPpred-positive;I=stringent intacelar-ike rule;O=remaining proteins.
Nand percentages were recalculated from the protein-level worksheets after accession-level deduplication.U is a prediction;I is not a measured contamination rate.AWF=apoplastic
Inclusion required≥2 unique peptides where available;root exudates used≥1.EV peptide counts were unavailable,so the reported EV list was retained without peptide-count filtering.

The comparison revealed two major patterns and distinct dominant protein classes across the nine datasets. First, unconventional candidates exceeded canonical candidates in both protein number and relative abundance in seven datasets, whereas suspension-cell culture medium showed the opposite pattern and root exudates exhibited similar proportions of the two predicted classes (Fig. 2J, K). Second, the combined proportion of canonical and unconventional candidates was higher in relative abundance than in protein number in seven datasets. Seed exudates and the cell-wall preparation were exceptions, where these two classes accounted for 51.6% versus 45.3% and 26.7% versus 14.9% of protein number and relative abundance, respectively (Fig. 2L). Finally, suspension-cell culture medium, the soluble cotton-fibre apoplastic fraction and the cell-wall protein preparation showed the highest proportions of canonical, unconventional and intracellular-like candidates, respectively. Canonical candidates accounted for 28.1% of proteins and 39.0% of relative abundance in suspension-cell culture medium, unconventional candidates for 40.3% and 47.2% in the soluble cotton-fibre apoplastic fraction, and intracellular-like proteins for 50.1% and 65.2% in the cell-wall preparation (Fig. 2J, K).

The proportion of unconventional candidates exceeded that of canonical candidates in both protein number and relative abundance in seven of the nine datasets. A similar pattern was reported in sugar beet leaf apoplastic fluid, where 38% of identified proteins were predicted to undergo unconventional secretion, compared with 26% assigned to canonical secretion (Ceballos-Laita et al., 2015). However, this relationship was not consistent across extracellular sample types. In oilseed rape xylem sap, 42% of identified proteins were classified as canonical secretory proteins and 25% as unconventional secretory proteins (Luo and Zhang, 2019), while a tomato xylem sap study reported 74% predicted canonical secretory proteins and 9% predicted leaderless secretory proteins (de Lamo et al., 2018). The suspension-cell culture medium likewise showed more canonical than unconventional candidates, whereas root exudates contained similar proportions of the two classes. Thus, although unconventional candidates predominated in seven datasets in our unified reanalysis, published results from comparable extracellular fractions and the suspension-culture sample show that this relationship is not universal. Differences in sample type, preparation and prediction criteria should be considered when interpreting these comparisons. Together, these comparisons suggest that the balance between predicted canonical and unconventional secretory protein classes reflects not only the biology of secretion but also how the sample was obtained.

Second, the combined proportion of canonical and unconventional candidates was higher in relative abundance than in protein number in seven datasets. This indicates that proteins assigned to these two predicted secretory classes had a higher mean relative abundance than proteins in the remaining classes within those fractions. Previous analyses of plant apoplastic fluid have also shown that protein-number and abundance-based profiles can differ substantially. In Arabidopsis leaf apoplastic fluid, proteins assigned to the apoplast accounted for 29% of identified proteins but 70.3% of the estimated abundance after adjustment for highly abundant Rubisco, whereas chloroplast-assigned proteins accounted for 26.7% of protein number but 15.5% of abundance (Karimi et al., 2025). A grapevine leaf apoplastic fluid study similarly found that pathogenesis-related proteins contributed 50.7% of the estimated protein content, illustrating how particular extracellular protein groups can dominate the abundance profile (Delaunois et al., 2013). These studies support the importance of considering protein abundance alongside protein number, although neither used our four-class prediction framework and therefore neither independently establishes that the two predicted secretion classes generally have a higher abundance proportion. In our comparison, seed exudates and the cell-wall preparation showed the opposite relationship, with the two predicted secretion classes accounting for 51.6% and 26.7% of protein number but only 45.3% and 14.9% of abundance, respectively. The direction of this difference therefore depends on the protein composition of the fraction analyzed and should not be treated as a universal feature of plant extracellular proteomes.

The highest proportions of canonical, unconventional and intracellular-like candidates were observed in suspension-cell culture medium, the soluble cotton-fibre apoplastic fraction and the cell-wall protein preparation, respectively. The high proportion of canonical candidates in suspension-cell culture medium is consistent with an independent study of Medicago suspension cultures, in which 34 of 39 identified extracellular-medium proteins contained predicted secretion signals, although this criterion is not identical to our canonical category (Kusumawati et al., 2008). The high proportion of unconventional candidates in the cotton-fibre dataset should be interpreted specifically for the soluble apoplastic fraction, as extracellular proteins differ in their association with the cell-wall matrix and may be recovered differently by extraction procedures (Rose and Lee, 2010). In contrast, the high proportion of intracellular-like proteins in the cell-wall preparation may partly reflect the co-extraction of intracellular proteins: an independent study showed that improved wall purification reduced predicted intracellular proteins and yielded a fraction in which 73% of identified proteins were predicted to be secreted (Feiz et al., 2006). Nevertheless, an intracellular-like prediction alone does not establish contamination.

Based on the target fractions, their preparation methods and the nine-dataset comparison, we propose a decision tree for selecting plant extracellular protein sampling and fractionation methods (Fig. 3).

**Fig. 3.**
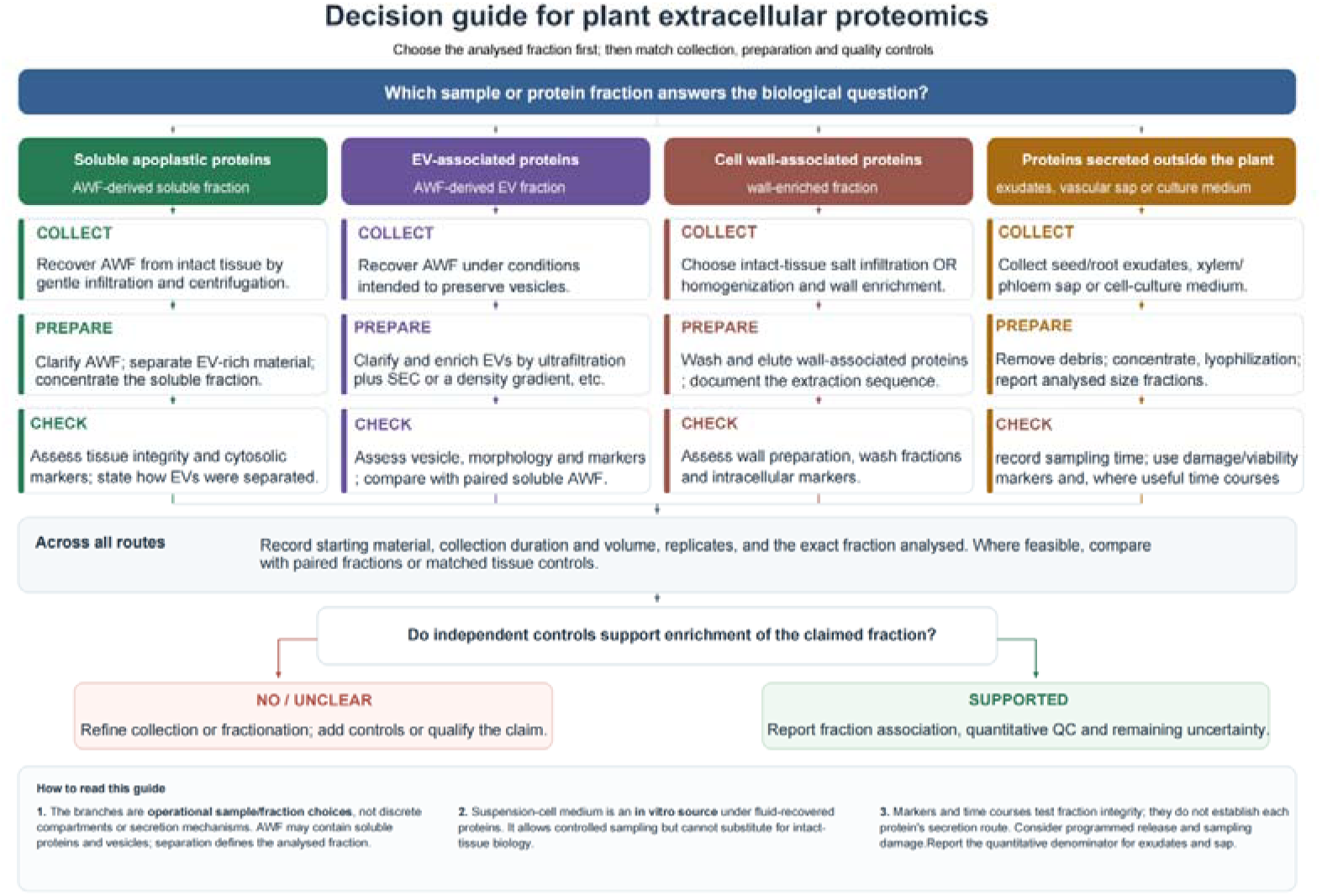
Decision tree for sampling and evaluating plant extracellular protein fractions. The research target determines one of four operational target fractions: (A) soluble apoplastic proteins, (B) extracellular vesicle (EV)-associated proteins, (C) cell wall-associated proteins, or (D) proteins recovered from exudates, vascular sap and culture medium. Each route links sample collection and fraction preparation to relevant quality checks. The final check asks whether the controls support enrichment of the intended fraction; weak support calls for better fractionation or a narrower interpretation. AWF, apoplastic washing fluid; SEC, size-exclusion chromatography; QC, quality control.

## 5. Establishing extracellular protein origin from proteomics data

Proteomics studies have employed several strategies to estimate whether detected proteins genuinely originate from the extracellular compartment. A straightforward strategy is comparative analysis against a whole-tissue or total-protein reference, in which proteins enriched in the extracellular fraction relative to the corresponding tissue extract are considered more likely to represent genuine extracellular components (Fig.4A, B). For example, Kourelis et al. (2019) classified proteins detected in *Nicotiana benthamiana* apoplastic fluid as apoplastic when they were either exclusively detected in the apoplastic fraction or strongly enriched relative to the total extract (log FC ≥ 1.5, p ≤ 0.01), while proteins enriched in the total extract were classified as intracellular. A second approach exploits the temporal dynamics of extracellular proteome collection (Fig. 4C, D). Proteins released through tissue disruption may be expected to be most abundant during the initial collection period, whereas proteins continuously present in the extracellular fluid may persist during subsequent sampling. Evidence for such temporal changes in xylem sap protein composition dates back to studies showing that individual protein isoforms can be restricted to the initial phase of stem exudation while others persist throughout collection (Biles and Abeles, 1991). Consecutive xylem sap fractions have also been analysed by proteomics, including our analysis of *Solanum tuberosum* and *Hordeum vulgare*, which demonstrated an enrichment of proteins annotated as extracellular in subsequent collection fractions (Kopecká and Černý, 2024). A third approach, and one that can extend these principles, is to examine protein abundance as a function of extraction severity (Fig. 4E, F). The underlying rationale is that increasing extraction force should have a progressively greater effect on proteins released through cellular disruption, whereas proteins already present in the extracellular compartment should be less dependent on extraction severity. Experimental evidence supports this premise: in maize roots, cytoplasmic contamination of apoplastic fluid was detectable after centrifugation even at 200 × g, whereas material collected using filter strips lacked detectable glucose-6-phosphate (Dragišić Maksimović et al., 2014). Similarly, recent proteomic analysis of *Arabidopsis* apoplastic fluid showed that the extraction procedure substantially influenced the resulting proteome and affected the extent of intracellular contamination (Chen, Karinshak and Vierstra, 2025b). These observations provide a rationale for treating extraction severity not only as a methodological variable but also as a potential analytical dimension. Systematic sampling across an extraction-severity gradient could therefore distinguish proteins whose abundance remains relatively stable from those showing a progressive increase with increasing extraction stress, providing an additional criterion for assessing extracellular versus leakage-derived protein populations.

**Fig. 4.**
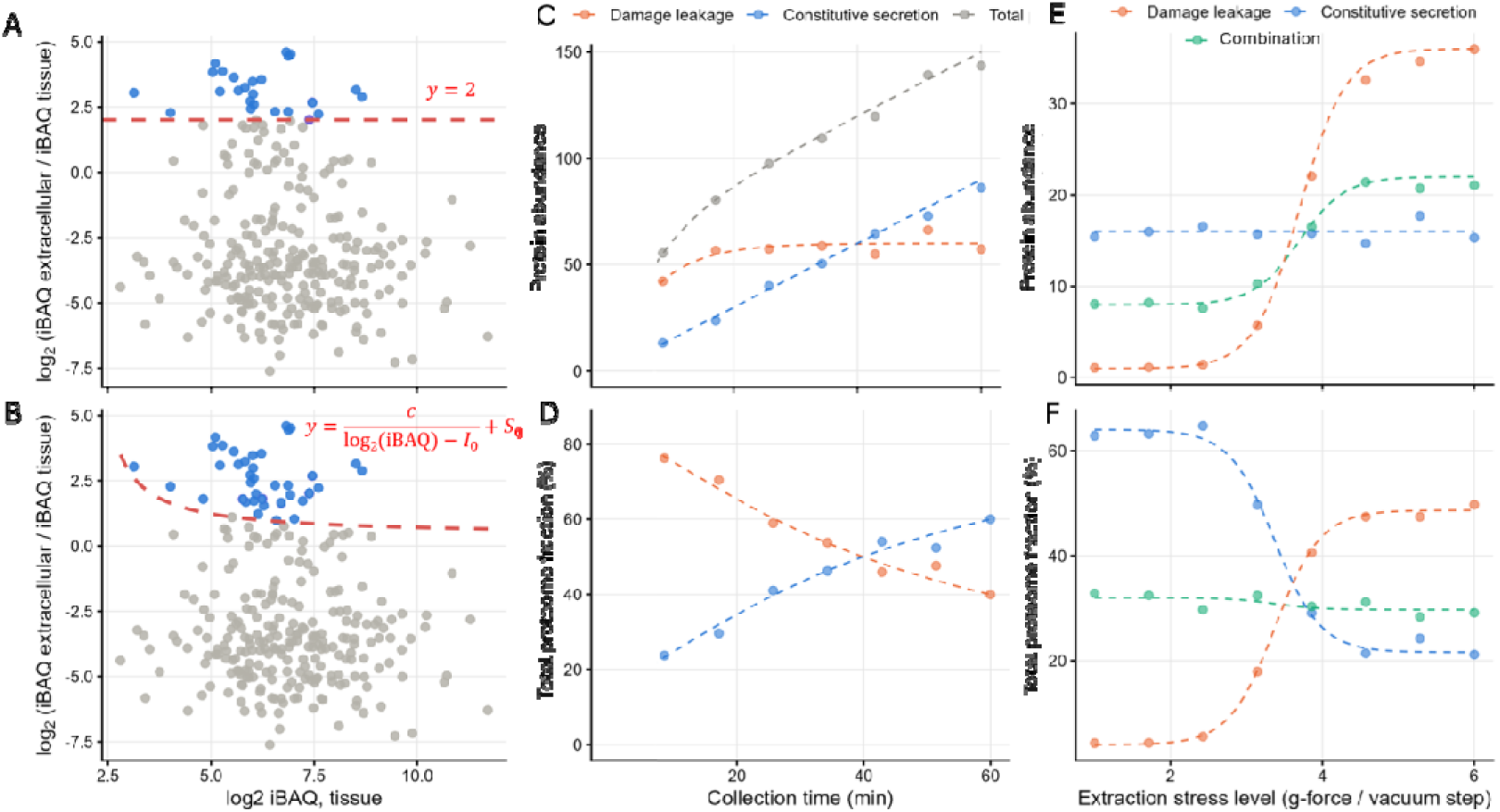
Estimating the extracellular origin of proteins from proteomics data. (A-B) Ratio-based classification using a complementary whole-tissue dataset. Enrichment is expressed as log2FC = log2(abundance in extracellular fraction / abundance in whole tissue). The rationale is that proteins abundant in whole tissue and not enriched in the extracellular fraction are more likely to be leakage contaminants than genuine extracellular components. (A) Standard classification with a single fixed log2FC threshold. (B) Adaptive classification in which the threshold depends on protein abundance. In this example, the adaptive threshold was applied with an abundance penalty factor of c = 2.5 and a lowered base threshold of log2FC > 0.4. Only proteins above a minimum abundance of I = 2 (log2iBAQ) were evaluated. (C, D) Kinetic classification from time-resolved sampling of a continuously collected fluid (e.g., exudate or xylem sap). Genuinely extracellular proteins accumulate throughout the collection period. Leaked proteins are released mainly at the earliest time points, after damage, and then plateau. (E, F) Classification by extraction-stress titration. Parallel samples are collected at increasing stress (higher centrifugal force, or higher vacuum for apoplastic fluid). Constitutively extracellular proteins remain relatively independent of stress level. Leaked proteins increase with stress as cell rupture increases. (D, F) Relative-abundance representations of the data shown in C and E. Because relative abundances are constrained by the total protein pool, accumulation of leaked intracellular proteins can cause genuine extracellular proteins to appear reduced even when their absolute abundance remains unchanged. This compositional bias should therefore be considered when interpreting relative proteomics data. All plotted values and thresholds are illustrative.

## 6. Challenges and future directions

### Determining the origin of extracellular proteins

Detection of a protein in an extracellular sample establishes its presence in the analyzed fraction, but does not show how it reached that fraction. Proteins may be secreted, released during normal development, or introduced through sampling-induced cell damage. These sources are particularly difficult to distinguish in seed and root exudates. Developmentally programmed cell death in the Arabidopsis root cap illustrates a possible source of released cellular contents, but does not establish the origin of any protein in the datasets analyzed here (Wang et al., 2024). Proteins may also have multiple localizations or appear in extracellular fractions through association with extracellular proteins or structures. Accordingly, the intracellular-like category in Table 2 is a prediction category, not a measured contamination rate. Assignment to the putative canonical or unconventional category likewise does not establish a specific secretion route (Robinson et al., 2016).

### Quality control of extracellular fractions

No single marker or threshold is suitable for every plant extracellular protein preparation. Quality control should reflect the target fraction. For leaf AWF, tissue integrity and the activity of a panel of intracellular marker enzymes should be assessed alongside extracellular enrichment; no single enzyme establishes contamination. Cell viability should also be monitored when collecting suspension-culture medium, because stress-induced loss of viability may contribute proteins from damaged cells (Ngcala et al., 2020). For cell wall preparations, studies should document wall enrichment and washing steps. EV preparations require particle characterization, protein markers, and appropriate controls. Comparing an EV-enriched fraction with a matched soluble apoplastic fraction can help determine whether a protein is enriched with vesicles (Rodríguez de Lope et al., 2025).

For exudates and vascular sap, sequential collection can help identify proteins potentially associated with sampling damage. In one xylem sap study, some proteins were abundant in the first collected fraction but declined rapidly thereafter, whereas proteins predicted to be extracellular were enriched in later fractions. The authors inferred that the early fraction was more affected by tissue damage at the cut surface (Kopecká and Černý, 2024). Guidance for plant EV studies recommends complementary particle and molecular characterization and complete reporting of isolation steps (Thieron et al., 2024; Welsh et al., 2024). Together, these measures assess enrichment and sample quality, but do not independently determine each protein’s route of release.

### Limitations of cross-study comparisons and localization prediction

The nine datasets analyzed here differ in species, tissue, developmental stage, treatment, sample preparation, and proteomics workflow. Protein counts and abundance profiles describe differences among the final analyzed fractions, but cannot isolate the effect of any single collection or purification step. Nor can the proportion assigned to the intracellular-like category be converted into an experimentally verified contamination rate. The “Other” category, comprising proteins not assigned to the canonical, unconventional or intracellular-like categories, is substantial in some datasets: it accounts for 58.5% of identified proteins in the suspension-culture dataset. Thus, the current classification does not resolve every protein.

Localization prediction helps organize candidates, but may miss proteins with multiple localizations or possible moonlighting functions, or proteins that enter an extracellular fraction through association with extracellular structures. Metabolic enzymes such as MDH, GAPDH and enolase illustrate this ambiguity: their activities are routinely used to estimate cytosolic contamination of apoplastic fluids (Alexandersson et al., 2013), yet MDH also occurs in the plant apoplast as a genuinely wall-bound enzyme rather than a cytoplasmic artefact (Hadži-Tašković Šukalović et al., 2011), and GAPDH is an established moonlighting protein with functions beyond the cytosol (Cao et al., 2025). Their detection in a proteome therefore warrants investigation but does not alone prove sampling damage. A recent benchmark of human protein localization predictors identified multiple localization as a continuing challenge; as its test set comprised human proteins, its performance estimates cannot be transferred directly to plants (Wefers et al., 2026). Predictions should therefore serve as supporting annotations, rather than as a quality-control gate in the method-selection decision tree or as proof of a protein’s extracellular localization.

### Practical priorities for future studies

A useful next step is to compare fractions prepared from the same biological material under clearly documented conditions. Retaining the starting sample and intermediate fractions, with suitable quantitative measurements, would help assess protein recovery and enrichment at each step. For exudates and vascular sap, studies should record collection time and, where feasible, collect sequential fractions. They should also report the amount of starting material, collection volume, protein recovery, and basis for abundance normalization. Normalization solely to total protein may obscure changes in the overall amount of protein released. Changes in protein composition should therefore be distinguished from changes in total release; comparison of absolute release amounts for individual proteins requires an appropriate quantitative standard.

These records and indicators of tissue damage are especially important when a stress treatment may alter both extracellular protein release and cell viability. Candidates of particular biological interest can then be examined using independent localization or functional experiments suited to the system. Such approaches would improve comparability across studies while preserving a necessary distinction: a well-characterized extracellular fraction supports an association between a protein and that fraction, but proteomic detection alone does not establish how the protein was released.

## Declaration of generative AI and AI-assisted technologies in the manuscript preparation process

During the preparation of this manuscript, the authors used Microsoft Copilot (M365 Copilot, GPT-5 based) to improve language and readability. After using this tool/service, the authors thoroughly reviewed and edited the content as needed and take full responsibility for the final published version of the article.

## References

Ødum, M.T., et al., 2024. DeepLoc 2.1: multi-label membrane protein type prediction using protein language models. Nucleic Acids Res, 52: W215–W220.

Albenne, C., Canut, H., Jamet, E., 2013. Plant cell wall proteomics: the leadership of Arabidopsis thaliana. Front Plant Sci, 4: 111.

Alexandersson, E., Ali, A., Resjö, S., Andreasson, E., 2013. Plant secretome proteomics. front plant sci 4:9

Almagro Armenteros, J.J., et al., 2019. Detecting sequence signals in targeting peptides using deep learning. Life Sci Alliance, 2: e201900429.

Bannai, H., et al., 2002. Extensive feature detection of N-terminal protein sorting signals. Bioinformatics, 18: 298–305.

Biles, C.L., Abeles, F.B., 1991. Xylem sap proteins. Plant Physiol, 96(2): 597–601.

Broussard, L., Abadie, C., Lalande, J., Limami, A.M., Lothier, J., Tcherkez, G., 2023. Phloem sap composition: what have we learnt from metabolomics? Int J Mol Sci, 24(8):6917

Cai, Q., Qiao, L., Wang, M., et al., 2018. Plants send small RNAs in extracellular vesicles to fungal pathogen to silence virulence genes. science 360, 1126–1129

Calderan-Rodrigues, M.J., et al., 2016. Cell wall proteome of sugarcane stems: comparison of a destructive and a non-destructive extraction method showed differences in glycoside hydrolases and peroxidases. BMC Plant Biol, 16: 14.

Calderan-Rodrigues, M.J., et al., 2019. Int J Mol Sci 20:1975

Cao, Y., Hong, J., Wang, H., Lin, M., Cai, Y., Liao, L., Li, X., Han, Y, 2025. Beyond glycolysis: multifunctional roles of glyceraldehyde-3-phosphate dehydrogenases in plants. Hortic Res, 12(6): uhaf070.

Carella, P., Wilson, D.C., Kempthorne, C.J., Cameron, R.K., 2016. Vascular sap proteomics: providing insight into long-distance signaling during stress. Front Plant Sci, 7: 651.

Ceballos-Laita, L., Gutierrez-Carbonell, E., Lattanzio, G., et al., 2015. Protein profile of Beta vulgaris leaf apoplastic fluid and changes induced by Fe deficiency and Fe resupply. Front Plant Sci, 6: 145.

Charura, N., Llamas, E., De Quattro, C., Vilchez, D., Nowack, M.K., Zuccaro, A, 2024. Root cap cell corpse clearance limits microbial colonization in Arabidopsis thaliana. elife 13, RP96266

Chen, H.-C., Newton, C.J., Diaz, G., et al., 2025a. Proteomic snapshot of pattern triggered immunity in the Arabidopsis leaf apoplast. Plant J, 123(6): e70498.

Chen, K.-E., Karinshak, M., Vierstra, R.D., 2025b. Streamline protocol for Arabidopsis apoplastic fluid isolation enables a detailed proteomic view of the plant extracellular space. Plant Direct, 9: e70087.

Chen, S., et al., 2022. Quantitative proteomics analysis of tomato root cell wall proteins in response to salt stress. frontiers in plant science 13, 1023388

Chou, K.-C., Shen, H.-B, 2010. Plant-mPLoc: A top-down strategy to augment the power for predicting plant protein subcellular localization. PLoS ONE, 5: e11335.

de Lamo, F.J., Constantin, M.E., Fresno, D.H., Boeren, S., Rep, M., Takken, FLW, 2018. Xylem sap proteomics reveals distinct differences between R gene- and endophyte-mediated resistance against Fusarium wilt disease in tomato. Front Microbiol, 9: 2977.

De Vallée, A., Dupuy, J.-W., Moriscot, C., et al., 2023. Extracellular vesicles of the plant pathogen Botrytis cinerea. journal of fungi 9, 495

De Palma, M., et al., 2020. Plant roots release small extracellular vesicles with antifungal activity. Plants, 9: 1777.

Delaunois, B., Colby, T., Belloy, N., et al., 2013. Large-scale proteomic analysis of the grapevine leaf apoplastic fluid reveals mainly stress-related proteins and cell wall modifying enzymes. BMC Plant Biol, 13(1): 24.

Ding, Y., Robinson, D.G., Jiang, L, 2014. Unconventional protein secretion (UPS) pathways in plants. current opinion in cell biology 29, 107–115

Domínguez, F., Cejudo, F.J., 2014. Programmed cell death (PCD): an essential process of cereal seed development and germination. Front Plant Sci, 5: 366.

Douché, T., et al., 2021. Cell wall proteomic datasets of stems and leaves of brachypodium distachyon. Data Brief, 35: 106818.

Dragišić Maksimović, J. J., Živanović, B., D., Maksimović, V. M., Mojović, M. D., Nikolić, M. T., Vučinić, Ž. B., 2014. Filter strip as a method of choice for apoplastic fluid extraction from maize roots. Plant Science, 223: 49–58.

Drula, E., et al., 2022. The carbohydrate-active enzyme database: functions and literature. Nucleic Acids Res, 50: D571–D577.

Eldahshoury, M.K., et al., 2024. Isolation of small extracellular vesicles from the apoplastic wash fluid of Nicotiana benthamiana leaves. Curr Protoc, 4: e70026.

Feiz, L., Irshad, M., Pont-Lezica, R.F., Canut, H., Jamet, E., 2006. Evaluation of cell wall preparations for proteomics: a new procedure for purifying cell walls from Arabidopsis hypocotyls. Plant Methods, 2: 10.

Gao, Y.Q., Su, Y., Chao, D.Y., 2024. Exploring the function of plant root diffusion barriers in sealing and shielding for environmental adaptation. Nat Plants, 10: 1865–1874.

Gao, J., Li, Y., Zhang, S., et al., 2026. Structural and functional characterization of EXPO-derived extracellular vesicles in plants. advanced science 13, e06163

Hadži-Tašković Šukalović, V., Vuletić, M., Marković, K., Vučinić, Ž, 2011. Cell wall-associated malate dehydrogenase activity from maize roots. Plant Science, 181(4): 465–470.

Hallgren, J., et al., 2022. DeepTMHMM predicts alpha and beta transmembrane proteins using deep neural networks. biorxiv [preprint]

Hooper, C.M., et al., 2022. Subcellular localisation database for Arabidopsis proteins version 5 [dataset]

Hooper, C.M., et al., 2020. CropPAL for discovering divergence in protein subcellular location in crops to support strategies for molecular crop breeding. Plant J, 104: 812–827.

Horton, P., et al., 2007. WoLF PSORT: protein localization predictor. Nucleic Acids Res, 35: W585–W587.

Huang, D., Chen, J., Zhao, M., Shen, H., Jin, Q., Xiao, D., Peng, Z., Chen, T., Zhang, Y., Rao, D., Liu, M., 2025. Plant-derived extracellular vesicles: composition, function and clinical potential. J Transl Med, 23: 1065

Huang, Y., Li, W., Liu, T., et al., 2025. Rice extracellular vesicles send defense proteins into fungus Rhizoctonia solani to reduce disease. developmental cell 60, 1168–1181.e6

Huang, Y., Wang, S., Cai, Q., Jin, H, 2021. Effective methods for isolation and purification of extracellular vesicles from plants. J Integr Plant Biol, 63: 2020–2030.

Jia, Y., Yu, L., Ma, T., et al., 2022. Small extracellular vesicles isolation and separation: current techniques, pending questions and clinical applications. Theranostics, 12(15): 6548.

Jian, Y., Liu, Z., He, P., et al., 2024. An emerging connected view: phytocytokines in regulating stomatal, apoplastic, and vascular immunity. Curr Opin Plant Biol, 82: 102623.

Jin, J., et al., 2022. PCMDB: A curated and comprehensive resource of plant cell markers. Nucleic Acids Res, 50: D1448–D1455.

Jung, Y.H., et al., 2008. J proteome res 7:5187

Kırbaş, O.K., Bozkurt, B.T., Asutay, A.B., Mat, B., Ozdemir, B., Öztürkoğlu, D., Ölmez, H., İşlek, Z., Şahin, F., Taşlı, P.N., 2019. Optimized isolation of extracellular vesicles from various organic sources using aqueous two-phase system. Sci Rep, 9(1): 19159.

Käll, L., Krogh, A., Sonnhammer, E.L.L., 2004. A combined transmembrane topology and signal peptide prediction method. J Mol Biol, 338: 1027–1036.

Karimi, H.Z., Chen, K.E., Karinshak, M., et al., 2025. Proteasomes accumulate in the plant apoplast where they participate in microbe-associated molecular pattern (MAMP)-triggered pathogen defense. Nat Commun, 16(1): 1634.

Koch, B.L., Rutter, B.D., Borniego, M.L., et al., 2025. Arabidopsis produces distinct subpopulations of extracellular vesicles that respond differentially to biotic stress, altering growth and infectivity of a fungal pathogen. journal of extracellular vesicles 14, e70090

Kolkas, H., et al., 2022. The cell wall proteome of Marchantia polymorpha reveals specificities compared to those of flowering plants. Front Plant Sci, 12: 765846.

Kopecká, R., Černý, M., 2024. Xylem sap proteome analysis provides insight into root-shoot communication in response to flg22. Plants, 13(14): 1983.

Kourelis, J., Kaschani, F., Grosse-Holz, F.M., Homma, F., Kaiser, M., van der Hoorn, R.A.L., 2019. A homology-guided, genome-based proteome for improved proteomics in the alloploid Nicotiana benthamiana. BMC Genomics, 20: 722.

Kranawetter, C., Sumner, L.W., 2025. Differential root zone secretions and the role of root border cells in rhizosphere manipulation. Phytochem Rev, 24(6): 5639–5658.

Kusumawati, L., Imin, N., Djordjevic, M.A., 2008. Characterization of the secretome of suspension cultures of Medicago species reveals proteins important for defense and development. J Proteome Res, 7(10): 4508–4520.

Lee, Y., Rani, H., Mallery, E.L., Szymanski, D.B., 2025. A cell fractionation and quantitative proteomics pipeline to enable functional analyses of cotton fiber development. Plant J, 121(4): e17246.

Liangsupree, T., Multia, E., Riekkola, M.L., 2021. Modern isolation and separation techniques for extracellular vesicles. J Chromatogr A, 1636: 461773.

Lonsdale, A., et al., 2023. LSPpred suite: tools for leaderless secretory protein prediction in plants. Plants, 12: 1428.

Luo, J.S., Zhang, Z., 2019. Proteomic changes in the xylem sap of Brassica napus under cadmium stress and functional validation. BMC Plant Biol, 19(1): 280.

McLaughlin, S., Himmighofen, P., Khan, S.A., Siffert, A., Robert, CAM, Sasse, J., 2025. Root exudation: an in-depth experimental guide. Plant Cell Environ, 48(5): 3052–3065.

Moreno-Sanguino, I., et al., 2026. Phloem proteomics to identify small open reading frame (sorf)-encoded peptides with a putative role in the control of flowering time in Arabidopsis. Physiol Plant, 178: e70860.

Nascimento, K.S., et al., 2018. Common features between the proteomes of floral and extrafloral nectar from the castor plant (Ricinus communis) and the proteomes of exudates from carnivorous plants. Front Plant Sci, 9: 549.

Neuberger, G., et al., 2003. Prediction of peroxisomal targeting signal 1 containing proteins from amino acid sequence. J Mol Biol, 328: 581–592.

Ngcala, M.G., et al., 2020. Heat stress triggers differential protein accumulation in the extracellular matrix of sorghum cell suspension cultures. Proteomes, 8: 29.

Pechanova, O., et al., 2010. Apoplast proteome reveals that extracellular matrix contributes to multistress response in poplar. BMC Genomics, 11: 674.

Perez-Riverol, Y., Bai, J., Bandla, C., et al., 2022. The PRIDE database resources in 2022: A hub for mass spectrometry-based proteomics evidences. Nucleic Acids Res, 50(D1): D543–D552.

Pernis, M., Danchenko, M., Jurcik, J., Kovac, A., Klubicová, K., 2026. Secretome profiling of Pinus nigra embryogenic cultures reveals a freely secreted protein fraction dominated by cell wall- and redox-related proteins. plant cell, tissue and organ culture 166, 86

Pope, R.E., Prade, R.A., 2026. Vesicle-driven endomembrane systems in fungi. Microbiol Mol Biol Rev, 90: e00297–24

Rejón, J.D., et al., 2013. Proteomics profiling reveals novel proteins and functions of the plant stigma exudate. J Exp Bot, 64: 5695–5705.

Robinson, D.G., Ding, Y., Jiang, L, 2016. Unconventional protein secretion in plants: A critical assessment. Protoplasma, 253: 31–43.

Rodríguez de Lope, M.M., Sánchez-Pajares, I.R., Herranz, E., et al., 2025. A compendium of bona fide reference markers for genuine plant extracellular vesicles and their degree of phylogenetic conservation. J Extracell Vesicles, 14(9): e70147.

Ropitaux, M., Bernard, S., Schapman, D., et al., 2020. Root border cells and mucilage secretions of soybean, Glycine max (merr) L.: characterization and role in interactions with the oomycete Phytophthora parasitica. cells 9, 2215

Rose, J.K., Lee, S.J., 2010. Straying off the highway: trafficking of secreted plant proteins and complexity in the plant cell wall proteome. Plant Physiol, 153(2): 433–6.

Routaboul, J.-M., et al., 2024. Arabidopsis hydathodes are sites of auxin accumulation and nutrient scavenging. the plant journal 120, 857–871

Rutter, B.D., Innes, R.W., 2017. Extracellular vesicles isolated from the leaf apoplast carry stress-response proteins. Plant Physiol, 173(1): 728–741.

Sánchez-López, C.M., et al., 2024. Phloem sap from melon plants contains extracellular vesicles that carry active proteasomes which increase in response to aphid infestation. J Extracell Vesicles, 13: e12517.

San Clemente, H., et al., 2022. Plant cell wall proteomes: the core of conserved protein families and the case of non-canonical proteins. Int J Mol Sci, 23: 4273.

Savojardo, C., et al., 2018. BUSCA: an integrative web server to predict subcellular localization of proteins. Nucleic Acids Res, 46: W459–W466.

Sigrist, C.J.A., et al., 2026. The PROSITE database for protein families, domains, and sites. Nucleic Acids Res, 54: D451–D458.

Silva, F.A., et al., 2020. LC-MS/MS based comparative proteomics of floral nectars reveal different mechanisms involved in floral defense of Nicotiana spp., Petunia hybrida and Datura stramonium. J Proteomics, 213: 103618.

Small, I., et al., 2004. Predotar: A tool for rapidly screening proteomes for N-terminal targeting sequences. Proteomics, 4: 1581–1590.

Sperschneider, J., et al., 2018. ApoplastP: prediction of effectors and plant proteins in the apoplast using machine learning. New Phytol, 217: 1764–1778.

Staudinger, C., et al., 2022. The wheat secreted root proteome: implications for phosphorus mobilisation and biotic interactions. J Proteomics, 252: 104450.

Strehmel, N., et al., 2017. Stress-related mitogen-activated protein kinases stimulate the accumulation of small molecules and proteins in Arabidopsis thaliana root exudates. Front Plant Sci, 8: 1292.

Teufel, F., et al., 2022. SignalP 6.0 predicts all five types of signal peptides using protein language models. nature biotechnology

Thieron, H., Krassini, L., Kwon, S., et al., 2024. Practical advice for extracellular vesicle isolation in plant-microbe interactions: concerns, considerations, and conclusions. J Extracell Vesicles, 13: e70022.

Tiddia, F., et al., 2023. The proteome of the imbibition spillage in Cicer arietinum L. current plant biology, 35-36, 100310

Toyofuku, M., Schild, S., Kaparakis-Liaskos, M., Eberl, L., 2023. Composition and functions of bacterial membrane vesicles. Nat Rev Microbiol, 21: 415–430.

Wang, J., Bollier, N., Andrade Buono, R., et al., 2024. A developmentally controlled cellular decompartmentalization process executes programmed cell death in the Arabidopsis root cap. Plant Cell, 36(4): 941–962.

Wang, X., Chung, K.P., Lin, W., Jiang, L., 2018. Protein secretion in plants: conventional and unconventional pathways and new techniques. J Exp Bot, 69(1): 21–37.

Wang, J., Ding, Y., Wang, J., et al., 2010. EXPO, an exocyst-positive organelle distinct from multivesicular endosomes and autophagosomes, mediates cytosol to cell wall exocytosis in Arabidopsis and tobacco cells. the plant cell 22, 4009–4030

Wefers, Z., Gupta, A., Ahmed, N., Zhang, X., Lundberg, E, 2026. A comprehensive benchmark of sequence-based subcellular localization predictors for human proteins. Nat Methods, 23(7): 1458–1469.

Welsh, J.A., Goberdhan, D.C.I., O’Driscoll, L., et al., 2024. Minimal information for studies of extracellular vesicles (MISEV2023): from basic to advanced approaches. J Extracell Vesicles, 13(2): e12404.

Weng, Y., Sui, Z., Shan, Y., Hu, Y., Chen, Y., Zhang, L., Zhang, Y., 2016. Effective isolation of exosomes with polyethylene glycol from cell culture supernatant for in-depth proteome profiling. Analyst, 141(15): 4640–6.

Wu, J., Lee, D.Y., Wang, Y., Kim, S.T., Baek, S., Kim, S.G., Kang, K.Y., 2014. Protein profiles secreted from phylloplane of rice leaves free from cytosolic proteins: application to study rice-Magnaporthe oryzae interactions. physiological and molecular plant pathology 88, 28–35

Yang, J., Wang, Y., Fu, Y., Wang, S., Luo, Y., Cui, X., Sun, C., Ji, J., Mo, J., Wang, Y., Liu, Y., Hu, X., Chen, X., Jia, Y., Luo, Y., 2025. Plant extracellular vesicles: a promising bionic medicine platform for disease treatment and drug delivery. Interdiscip Med, 3: e20240101

Yugay, Y., et al., 2023. Isolation and characterization of extracellular vesicles from Arabidopsis thaliana cell culture and investigation of the specificities of their biogenesis. Plants, 12: 3604.

Zhao, L., et al., 2019. OutCyte: A novel tool for predicting unconventional protein secretion. Sci Rep, 9: 19448.

Zheng, L., Liu, Q., Wu, R., et al., 2023. The alteration of proteins and metabolites in leaf apoplast and the related gene expression associated with the adaptation of Ammopiptanthus mongolicus to winter freezing stress. Int J Biol Macromol, 240: 124479.

Zheng, L., Li, B., Liu, Q., Richardson, J.E., Zhou, Y., Gao, F., 2026. Proteomic analysis of leaf apoplast reveals that a jasmonate-regulated CHIA participates in the response to cold and drought stress in jojoba. Hortic Plant J, 12(7): 1697–1718.

